# G3DC: a Gene-Graph-Guided selective Deep Clustering method for single cell RNA-seq data

**DOI:** 10.1101/2023.01.15.524109

**Authors:** Shuqing He, Jicong Fan, Tianwei Yu

## Abstract

Single-Cell RNA sequencing (scRNA-seq) technology measures the expression of thousands of genes at the cellular level. Analyzing single cell transcriptome allows the identification of heterogeneous cell groups, cellular-level regulations, and the trajectory of cell development. An important aspect in the analyses of scRNA-seq data is the clustering of cells, which is hampered by issues such as high dimensionality, cell type imbalance, redundancy, and dropout. Given cells of each type are functionally consistent, incorporating biological relations between genes may improve the clustering results. Here, we develop a deep embedded clustering method, G3DC, that incorporates a graph loss based on existing gene network, together with a reconstruction loss to achieve both discriminative and informative embedding. The involvement of the gene network strengthens clustering performance, while helping the selection of functionally coherent genes that contribute to the clustering results. In addition, this method is well adapted to the sparse and zero-inflated scRNA-seq data with the ***ℓ*2**,**1**-norm involved. Extensive experiments have shown that G3DC offers high clustering accuracy with regard to agreement with true cell types, outperforming other leading single-cell clustering methods. In addition, G3DC selects biologically relevant genes that contribute to the clustering, providing insight into biological functionality that differentiate cell groups.

## 1 Introduction

The single-Cell RNA sequencing (scRNA-seq) technology enables precise descriptions of a tissue’s molecular states such as cell composition and gene expression characteristics of different cell types. Unsupervised clustering analysis is used to identify cell groups and provides meaningful explanations for gene expression variations. Recently, a number of clustering methods have been developed for clustering cells into different groups for better data interpretation and medical applications.

There exist traditional clustering methods such as K-means and hierarchical clustering. K-means [1] is based on the assumption that clusters can be well described by the centers, which may not be satisfied in real applications. Recently, a few variants of K-means have been proposed. For instance, pcaReduce [2] is a clustering method based on PCA [3], K-means and iterative hierarchical clustering. Starting from big clusters, it iteratively combines similar clusters and removes the principal component that explains the lowest degree of freedom after each iteration. RaceID3 [4] is designed for rare cell types and the method treats those rare cells as outliers in the regular clustering methods. To address the problem of using k-means with large datasets, mbkmeans [5] implements a mini-batch k-means algorithm and provides fast and scalable clustering of scRNA-seq data. Hierarchical clustering does not require the knowledge of cluster numbers and is used as one of the steps in many scRNA-seq clustering analyses. CIDR [6] is a fast and accurate clustering method through imputation and dimension reduction. It utilizes an implicit imputation approach to circumvent the dropout effects. BackSPIN [7] utilizes a deterministic biclustering method based on hierarchical clustering. The distinct subclasses of cells can be identified through repeated biclustering. SINCERA [8] identifies cell types using hierarchical clustering with similarity measurement based on centered Pearson’s correlation and average linkage.

Spectral clustering [9] is based on connectivity and originated from the graph partition problem [10]. Spectral clustering aims at finding the best partition to lower the weights of edges between different groups as much as possible. Louvain [11] and Leiden [12], which are used in single-cell RNAseq analysis tools like Seurat [13] and Scanpy [14], also partition the gene graph by iteratively aggregating nodes. Louvain adapts modularity optimization and community aggregation process, while the Levian algorithm adds a second phase of the refinement of partition in between [15].

Some other graph-based clustering methods were developed to capture the characteristics of the gene-gene expression relations. SNN-Clip (shared nearest neighbor) is a quasi-clique-based clustering algorithm to identify similar nodes that belong to the same cluster [16]. SIMLR learns a sample-to-sample similarity from the expression matrix by combining multiple Gaussian kernels [17]. Seurat measures global gene expression and separates cells from native spatial context [13]. Other clustering methods inlcude density-based methods like monocle2 [18], which can perform differential expression analysis between subpopulations of genes.

Cluster ensemble is a strategy to combine different clustering methods to enhance performance. Methods like SAFE [19], SAME [20], Sc-GPE [21] and scConsensus [22] are ensemble clustering algorithms based on and combine clustering methods like SC3, CIDR, SIMLR, SSNN-Louvain [23], MGPS-Louvain [24] and SNN-Clip. SC3 [25] overcomes the K-means problem of sensitivity to outliers by using multiple clustering solutions to perform the consensus clustering. And it has shown high accuracy and beat most of the other clustering methods. Based on SC3, scConsensus [22] obtains a consensus set of clusters based on at least two different clustering results. One common limitation of SC3 and scConsensus is that they have high computational complexity, which restricts their application to large scRNA-seq datasets. In addition, SC3 is based on multiple heuristic data preprocessing steps such as eliminating the low-expression or high-expression genes to achieve lower dimensions, which require strong prior knowledge.

In recent years, deep learning methods begin to play important roles in single-cell clustering problems. DCA [26] (deep count encoder) uses autoencoder to denoise scRNA-seq data and utilizes a zero-inflated negative binomial (ZINB) loss function. Tian et al. [27] proposed a model-based deep embedded clustering method with running time increasing linearly with sample size. DESC [28], based on DEC [29], first pretrains an autoencoder to learn a lowdimensional representation of input data and initialize the clusters, and then iteratively refines the clusters using the encoder and an auxiliary target distribution derived from the current soft cluster assignment. scCAN [30] combines autoencoder and network fusion and spectral clustering for cell segregation. It also uses a combination of the network fusion approach and k-NN algorithm to cluster big data. Considering the underlying relationships among cells as a graph structure, sigDGCNb [31] combines a graph convolutional network (GCN) [32] and an unsupervised deep clustering method to form the target distribution.

The existing methods still cannot address all the challenges of single-cell clustering. The first one is that the dimension of features (number of genes) is usually very high, making it really hard to directly perform clustering on the original data. Second, although heuristic gene/feature selection or other preprocessing methods can be used, the preserved genes or extracted features are not guaranteed to be discriminative and hence may lead to low clustering performance. Finally, current methods only focus on learning the embedded representation of genes but ignore the functional relationship between genes. Utilizing functional relationships between genes could potentially make cell clusters more functionally consistent. Therefore, in this work, we try to address these issues via clustering with adaptive feature selection, guided by the existing gene graph.

We propose a gene-graph-guided selective deep clustering method: G3DC, which is able to cluster cells and at the same time select important genes for further functional analysis. G3DC follows the idea of iDEC [29], which optimizes the latent representation and cluster assignment simultaneously. G3DC incorporates the gene interaction graph and encourage adjacent genes to share similar weights in a neural network. While strengthening the clustering performance, G3DC achieves the effect of group-sparse regularization that tends to select genes on subgraphs, making downstream interpretation easy. Further interpretations can be made to find which biological processes contribute to the separation of the clusters to shed light on the functional aspects in the cell clusters.

## 2 Results

### 2.1 Method overview

An overview of G3DC is shown in Figure 1. G3DC has four components. The first component is an autoencoder, which aims to learn a low-dimensional representation of the input high-dimensional data. The second component of G3DC is a clustering module, which is based on the KL divergence between a pseudo target distribution and a distribution given by the neural network. The third component is the feature selection regularizer, which is based on the *ℓ*_21_-norm performed on the weight matrix of the input layer of the autoencoder. The last component of G3DC is the gene graph module, which directly guides the feature selection and indirectly guides the clustering using the structural information between genes. Thus, the optimization objective of G3DC is composed of four parts: a reconstruction loss of the autoencoder, a clustering loss based on the KL divergence, a feature selection term based on the *ℓ*_21_ norm regularization, and a gene graph guidance term. G3DC utilizes the Laplacian matrix of the gene-gene interaction graph to make adjacent genes have similar weights, and hence guides the feature selection, reconstruction, and clustering. Importantly, the clustering-oriented and graph-guided objective enables G3DC to select informative and discriminative genes, which addresses the high dimensionality and sparsity of scRNA-seq data.

**Fig. 1.**
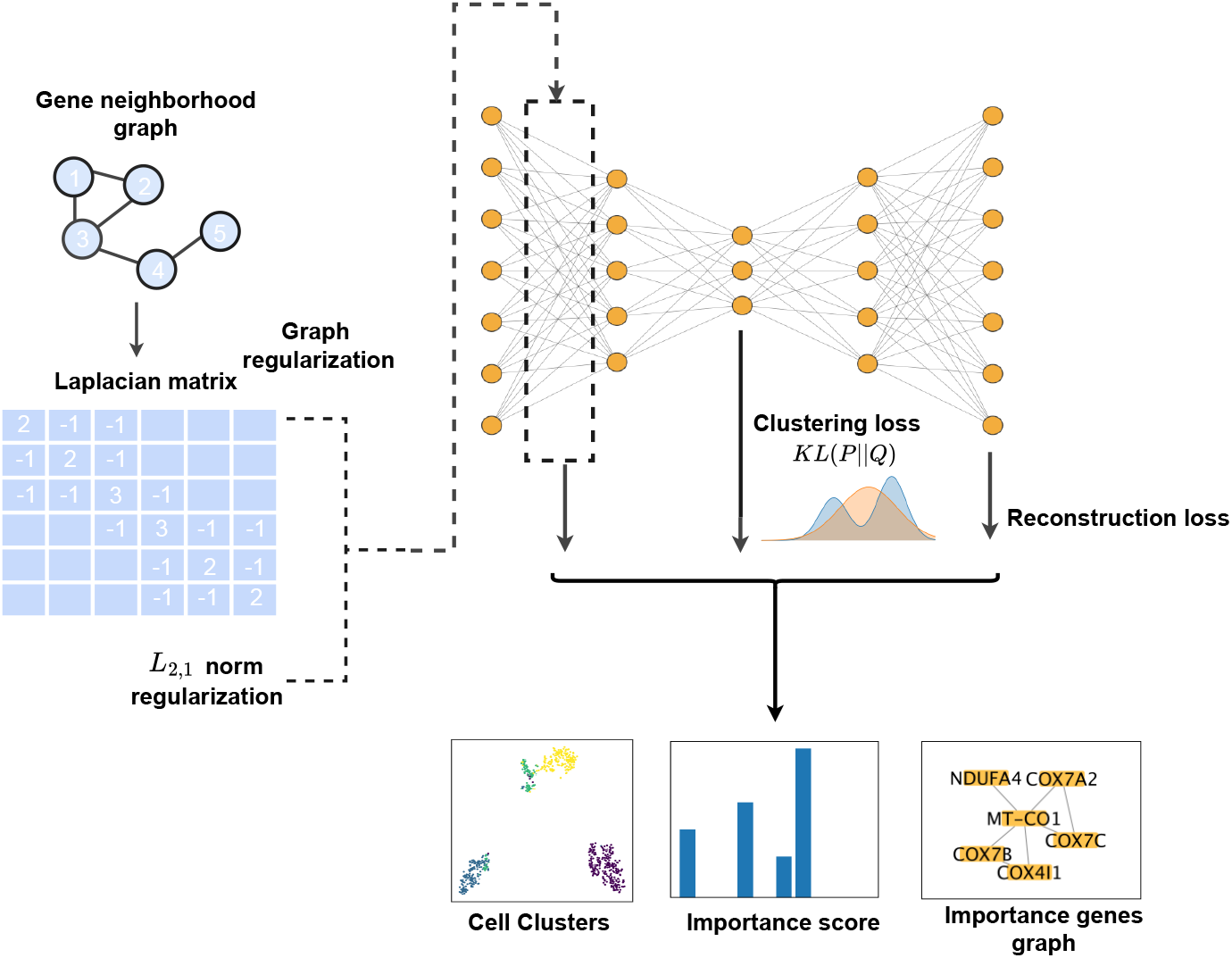
An overview of our proposed method. G3DC consists of an autoencoder, a clustering module, a feature selection module, and a gene graph module. The hidden layer of the autoencoder is used to learn the embedded feature representation, on which clustering is carried out. The KL divergence is computed on the hidden layer optimizing the similarity between a target distribution P and a learned distribution Q. The total optimization function is composed of four parts: 1) the clustering loss based on the KL divergence; 2) the reconstruction loss of the autoencoder; 3) the *ℓ*_2,1_-norm regularization for gene selection; 4) the gene graph regularization term Tr(*W*_1_*LW*_1_^⊤^).

### 2.2 Application to real single-cell data

G3DC was tested on five real scRNA-seq datasets to evaluate its performance (Table 1). Three of the datasets were subjected to variation-based gene screening, while the other two were of original dimension. The purpose was to test our method against datasets with different number of variables and ensure its robustness. The gene interaction networks were constructed using interactomes from the HINT database [33]. Detailed description of the datasets can be found in the Real datasets section.

**Table 1.**
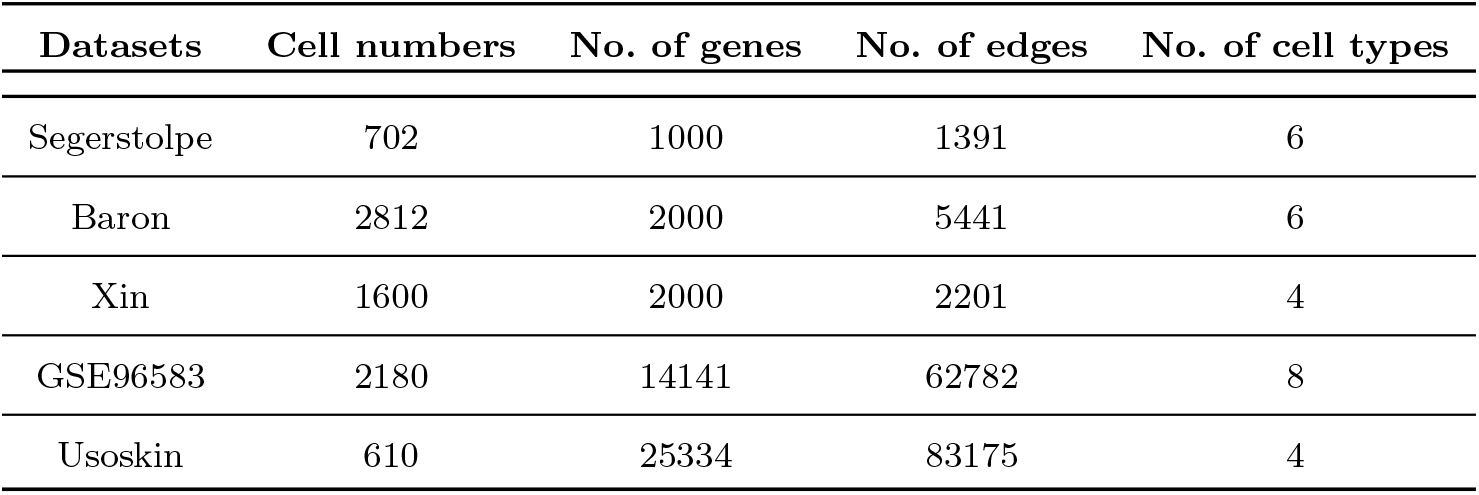
Summary of the real scARNA-seq datasets

To ensure all the genes in each dataset have similar scales, we performed the min-max normalization such that the expression values of each gene is in the range [0, 1]. This process ensures that the weights of the neural network for different genes are comparable and hence can be indicators for the selection of genes that contribute the most to the clustering results.

### 2.3 Comparison with other unsupervised clustering methods

G3DC was compared with three other methods of clustering: K-means [1], SC3 [25], and scDeepCluster [27]. K-means is a well-known traditional clustering method. SC3 has been one of the most competitive methods in single-cell clustering. scDeepCluster is a representative deep learning clustering method for scRNA-seq data. It learns a latent embedded representation that is optimized for clustering high-dimensional input in a non-linear manner.

After performing clustering, three evaluation metrics were used to compare the similarity between the clustering result and the true cell labels. They were: CA (Clustering Accuracy), NMI (Normalized Mutual Information), and ARI (Adjusted Rand Index). Detailed description about the three metrics is provided in the Evaluation metrics section. For each dataset, all methods were tested 20 times to check their accuracy and stability.

Figure 2 presents the overall performance of the four methods on the aforementioned five datasets. The distribution of the metrics from 20 repeated trials are shown on the graph using box plots. As indicated by the box plots, G3DC has high clustering stability as the range of the boxes is relatively small compared to other methods. The results showed that G3DC outperformed Kmeans and scDeepCluster on all the five datasets. It performed better than SC3 on Segerstolpe, Baron and GSE96583 data, while it was slightly defeated by SC3 on Baron data. G3DC’s CA and ARI exceeded that of SC3 on Usoskin data. The results indicate G3DC is highly competitive against other methods, including one of the leading methods SC3.

**Fig. 2.**
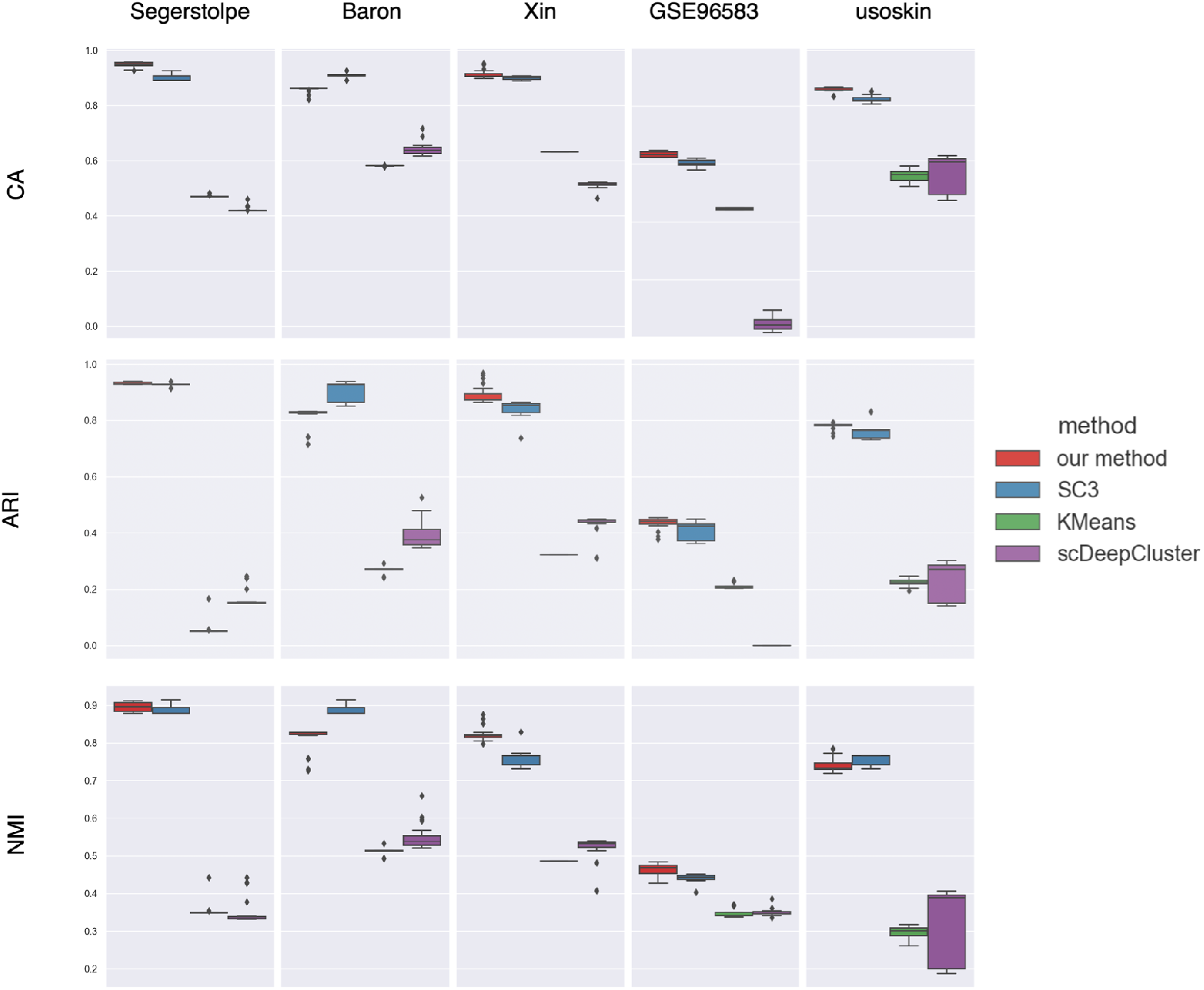
Comparison of clustering performances.

The hidden space in the middle of the autoencoder represents the lowdimensional embedded representation of the input gene expression data. To visualize the effectiveness of the deep embedding, three datasets were selected and t-SNE [34] was adopted to visualize those embedded points in a 2-D plane. The embedded points were extracted from the hidden layer of our model. Given K-means and SC3 are not suitable for this comparison, three other methods, vanilla deep embedding clustering (DEC), PCA, and scDeepCluster were selected to compare the embedding performance. Figure 3 shows that G3DC separated Baron and Segerstolpe data clearly into six types. Although vanilla DEC and PCA both achieved competitive 2D visualization effects, they wrongly mixed some small clusters into one, making the t-SNE visualization look like 3-4 clusters while indeed there were six. In contrast, G3DC successfully revealed the minor cell types and separated them from others. For scDeepCLuster, it seems to cluster less tightly on the Baron and Segerstolpe datasets. Similar results were observed on the Usoskin dataset. G3DC turns out to separate different cell types most effectively with only a few outliers compared to other methods in the low-dimensional embedded representation.

**Fig. 3.**
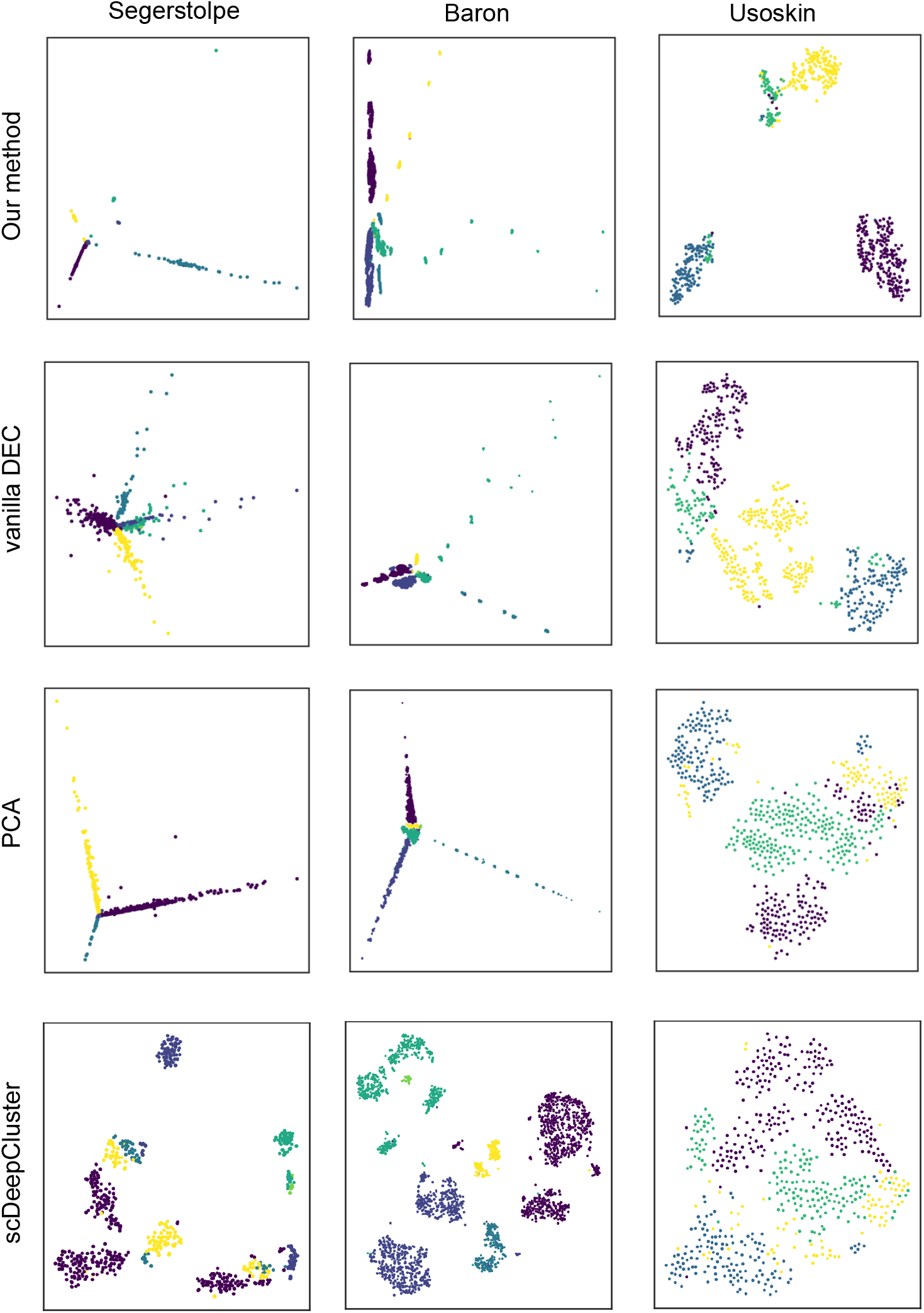
Comparison of 2D visualization of embedded representations

### 2.4 Example gene selection results

#### 2.4.1 Segerstolpe dataset

The Segerstolpe dataset was subjected to pre-screening, and only the top 1000 genes with highest variation levels entered the study. Based on our method of estimating variable importance, we selected the top 200 genes. We display those important genes having at least one connections between them in Figure 4. There were a total of 87 edges between 77 of the selected genes, and 68 selected genes formed a connected component of the network. Given the Segerstolpe dataset we used was a sub-dataset with only 1000 genes, the method showed a strong tendency to select connected genes on the input gene graph, making the selection functionally consistent. The results indicate that the trace penalty based on the input gene graph was effective.

**Fig. 4.**
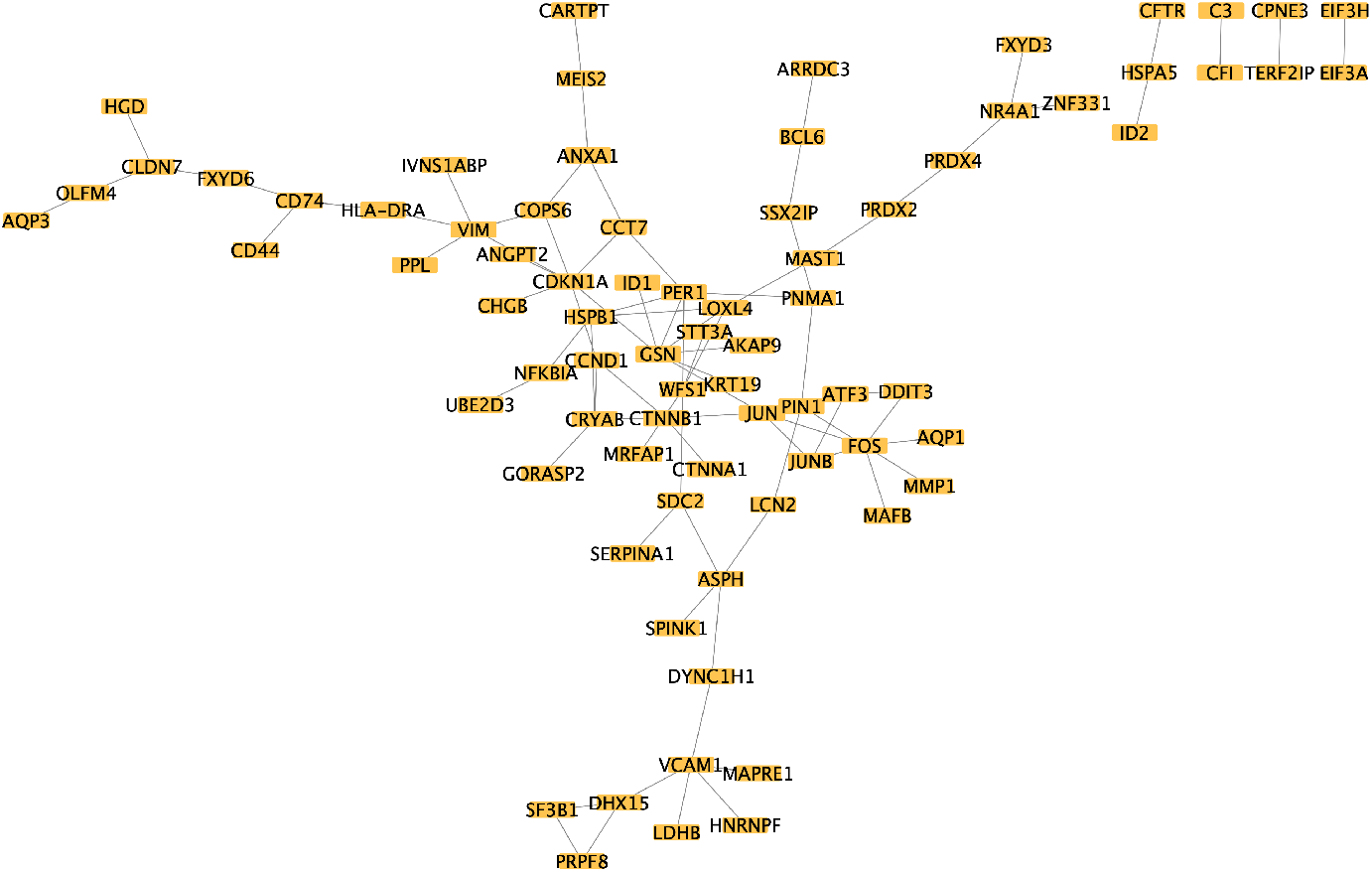
Connected graph of selected important genes in Segerstolpe pancreas dataset. Yellow nodes represent all the selected important genes that have at least one connection.

We further analyzed the selected genes using gene set enrichment analysis by GOstat [35]. To remove redundancy in the selected GO terms, we first deleted the GO terms with more than 200 genes. Then we look at all the pairwise relationships of the most significant GO terms. Each time we compare two GO terms and identify their shared genes. If the shared genes made up more than 75% of the genes in the larger GO term, we deleted the larger GO term. Else if the shared genes made up more than 90% of the smaller GO term, we deleted the smaller one. The top 15 GO terms are shown in Table 2.

**Table 2.** Top significant GO terms of Segerstolpe pancreas dataset.

| GOBPID | Term | Pvalue | OddsRatio | ExpCount | Count | Size |
| --- | --- | --- | --- | --- | --- | --- |
| GO:0031667 | response to nutrient levels | 1.23E-07 | 5.385802469 | 9.298342541 | 25 | 45 |
| GO:0071310 | cellular response to organic substance | 3.26E-07 | 2.59295393 | 38.0198895 | 64 | 184 |
| GO:0009968 | negative regulation of signal transduction | 5.13E-07 | 3.03962704 | 22.72928177 | 44 | 110 |
| GO:0009605 | response to external stimulus | 5.21E-07 | 2.501844784 | 40.91270718 | 67 | 198 |
| GO:0009628 | response to abiotic stimulus | 1.01E-06 | 3.011494253 | 21.6961326 | 42 | 105 |
| GO:0033554 | cellular response to stress | 2.47E-06 | 2.538619275 | 31.61436464 | 54 | 153 |
| GO:0061061 | muscle structure development | 1.65E-05 | 3.53909465 | 11.36464088 | 25 | 55 |
| GO:0009612 | response to mechanical stimulus | 2.35E-05 | 6.375080283 | 4.752486188 | 14 | 23 |
| GO:0007166 | cell surface receptor signaling pathway | 4.53E-05 | 2.255474453 | 30.99447514 | 50 | 150 |
| GO:0051592 | response to calcium ion | 4.62E-05 | 5.729479769 | 4.959116022 | 14 | 24 |
| GO:0014070 | response to organic cyclic compound | 4.97E-05 | 2.620235935 | 19.21657459 | 35 | 93 |
| GO:0009314 | response to radiation | 5.77E-05 | 4.705096074 | 6.198895028 | 16 | 30 |
| GO:0071396 | cellular response to lipid | 7.86E-05 | 3.506746988 | 9.504972376 | 21 | 46 |
| GO:0043408 | regulation of MAPK cascade | 0.00022625 | 2.873269062 | 12.19116022 | 24 | 59 |
| GO:0006974 | cellular response to DNA damage stimulus | 0.000241431 | 3.678947368 | 7.438674033 | 17 | 36 |

The biological processes that contribute the most to the clustering results were clearly associated with the function of the pancreas, with the majority of the top GO terms belonging to response to various nutrient conditions. For example, 64 of the selected genes fall into the process “cellular response to organic substance”. At the same time, a large portion of the genes were selected from signal transduction processes, such as “cell surface receptor signaling pathway” and “regulation of MAPK cascade”, signifying their importance in the regulation in pancreas function and nutrient response. In recent years, the MAPK (mitogen-activated protein kinases) pathways have been extensively studied in the context of pancreas cancer [36]. In regular cellular functions, MAPK signalling pathway was found to be responsible for the glucose-dependent activation of insulin gene promoters [37], and the MAPK pathway is implicated in type-1 and type-2 diabetes [38, 39].

#### 2.4.2 Usoskin data

The Usoskin dataset contains 25334 genes. Thus we selected the top 400 genes. We display the important genes having at least one connection in the input network in Figure 5. There were 115 such important genes with 131 edges, and 77 of them are involved in the largest component.

**Fig. 5.**
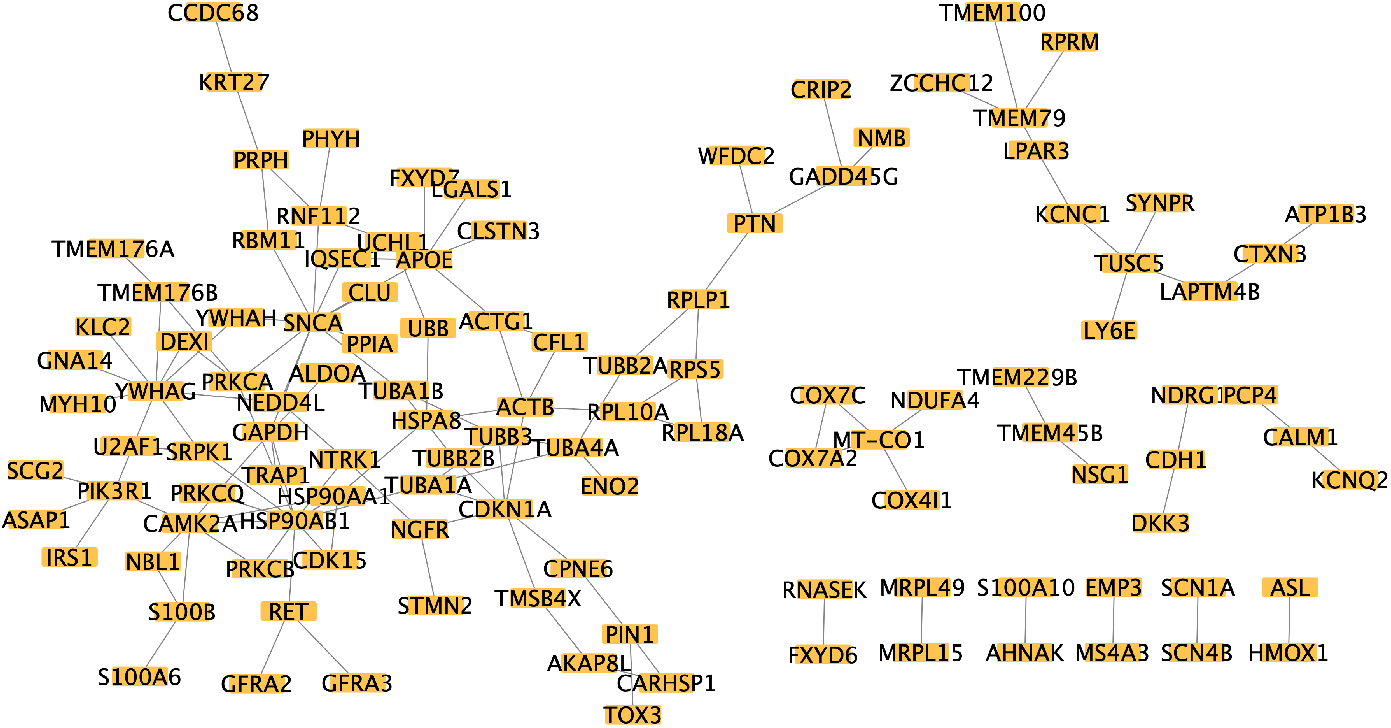
Connected graph of selected important genes in the Usoskin dataset. Yellow nodes represent all the selected important genes that have at least one connection.

We further analyzed the selected genes using gene set enrichment analysis, and removed redundancy following the same procedure as in the previous subsection. The top 15 biological processes are shown in Table 3. The Usoskin dataset contained measurement of four types of sensory neuron cells. It is clear that the top biological processes are mostly surrounding three general themes: (1) neural organization and structure, *e*.*g*. “postsynapse organization”, “synapse assembly”, “cell junction maintenance” *etc*; (2) sensory functions, *e*.*g*. “response to temperature stimulus”, “response to pain”, “response to heat” *etc*; (3) ion channel activity, *e*.*g*. “regulation of cation channel activity”, “positive regulation of cation transmembrane transport”, “regulation of voltage-gated calcium channel activity”, *etc*. Clearly the genes that differentiate the cell clusters were functionally meaningful based on the types of cells under study.

**Table 3.** Top significant GO terms of the Usoskin dataset.

| GOBPID | Term | Pvalue | OddsRatio | ExpCount | Count | Size |
| --- | --- | --- | --- | --- | --- | --- |
| GO:0009266 | response to temperature stimulus | 1.26E-07 | 6.620469083 | 2.382181646 | 14 | 161 |
| GO:0099173 | postsynapse organization | 2.40E-07 | 5.771928735 | 2.900047222 | 15 | 196 |
| GO:0048265 | response to pain | 1.48E-06 | 14.91079545 | 0.577050212 | 7 | 39 |
| GO:0016048 | detection of temperature stimulus | 1.79E-06 | 20.38804348 | 0.384700142 | 6 | 26 |
| GO:0023041 | neuronal signal transduction | 1.27E-05 | 201.8924731 | 0.059184637 | 3 | 4 |
| GO:0043954 | cellular component maintenance | 3.93E-05 | 8.509545455 | 0.932158036 | 7 | 63 |
| GO:2001257 | regulation of cation channel activity | 4.99E-05 | 4.722947417 | 2.530143239 | 11 | 171 |
| GO:0007416 | synapse assembly | 6.82E-05 | 4.550771351 | 2.618920195 | 11 | 177 |
| GO:0009408 | response to heat | 7.71E-05 | 6.34561195 | 1.390838974 | 8 | 94 |
| GO:0042982 | amyloid precursor protein metabolic process | 8.54E-05 | 7.442670455 | 1.05052731 | 7 | 71 |
| GO:1904064 | positive regulation of cation transmembrane transport | 0.000118834 | 4.659363745 | 2.322997009 | 10 | 157 |
| GO:2001259 | positive regulation of cation channel activity | 0.000143173 | 6.802545455 | 1.139304266 | 7 | 77 |
| GO:0034331 | cell junction maintenance | 0.000177216 | 10.91533714 | 0.532661735 | 5 | 36 |
| GO:1901385 | regulation of voltage-gated calcium channel activity | 0.000177216 | 10.91533714 | 0.532661735 | 5 | 36 |
| GO:0051651 | maintenance of location in cell | 0.000201351 | 3.992034206 | 2.959231859 | 11 | 200 |

#### 2.4.3 GSE96583 data

The GSE96583 dataset contains 14141 genes. We selected the top 400 genes with the greatest *ℓ*_2,1_-norm of the first network layer weights. Among them, 346 genes were connected with other selected genes, and 339 genes form a connected component. We display the genes having at least one connections between them in Figure 6.

**Fig. 6.**
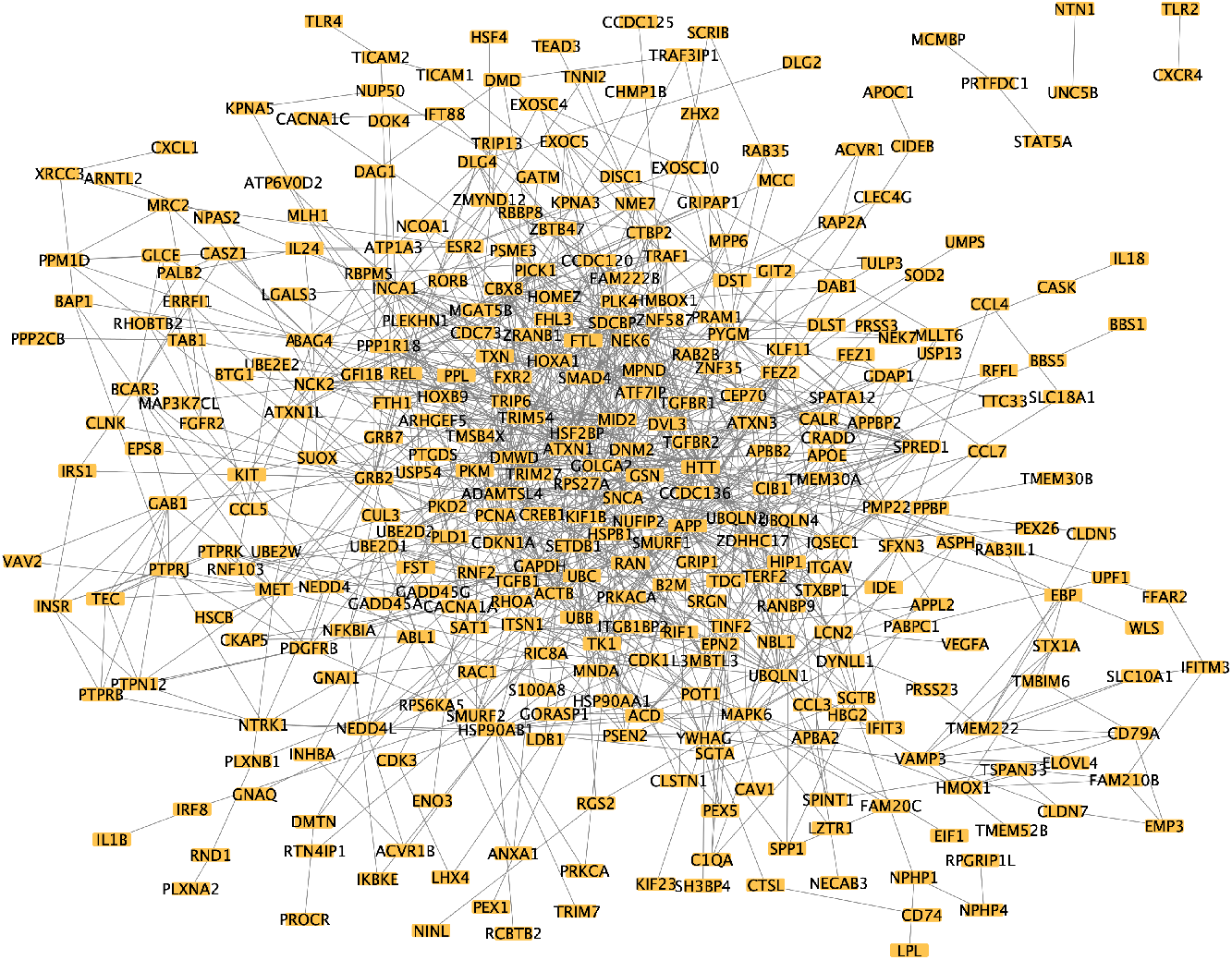
Connected graph of selected important genes in the GSE96583 dataset. Yellow nodes represent all the selected important genes that have at least one connection.

Functional analysis of the selected genes was conducted using GOstat (Table 4). The GSE96583 dataset studies 8 types of peripheral blood mononuclear cells (PBMC). The top biological processes were clearly consistent with the cell population. Most of the top biological processes belong to several categories that are intertwined with each other: (1) Cytokine response and chemotaxis, *e*.*g*. “regulation of natural killer cell chemotaxis”, “neutrophil chemotaxis”, “response to interferon-gamma”, *etc*; (2) Signalling pathways, *e*.*g*. “positive regulation of I-kappaB kinase/NF-kappaB signaling”, “protein kinase B signaling”, “chemokine-mediated signaling pathway”, *etc*; (3) Other cellular functions closely related to cell activation, *e*.*g*. “positive regulation of calcium ion transport”, “positive regulation of nitric oxide biosynthetic process” [40], “regulation of protein binding” *etc*. Overall, the results clearly pointed to some plausible biological processes and pathways that differentiate different types of PBMC cells.

**Table 4.** Top significant GO terms of GSE96583 dataset.

| GOBPID | Term | Pvalue | OddsRatio | ExpCount | Count | Size |
| --- | --- | --- | --- | --- | --- | --- |
| GO:2000501 | regulation of natural killer cell chemotaxis | 8.32E-11 | 250.8631579 | 0.097476066 | 6 | 8 |
| GO:0030593 | neutrophil chemotaxis | 3.53E-09 | 11.57865169 | 1.230635335 | 12 | 101 |
| GO:0070098 | chemokine-mediated signaling pathway | 8.29E-09 | 12.37226174 | 1.060052219 | 11 | 87 |
| GO:0034341 | response to interferon-gamma | 1.14E-08 | 9.10617975 | 1.657093124 | 13 | 136 |
| GO:0051928 | positive regulation of calcium ion transport | 1.31E-07 | 9.203391627 | 1.376849434 | 11 | 113 |
| GO:0045429 | positive regulation of nitric oxide biosynthetic process | 3.28E-07 | 18.9474313 | 0.463011314 | 7 | 38 |
| GO:0009266 | response to temperature stimulus | 1.17E-06 | 6.493670886 | 2.071366406 | 12 | 170 |
| GO:0050792 | regulation of viral process | 2.12E-06 | 6.786995691 | 1.815491732 | 11 | 149 |
| GO:0043123 | positive regulation of I-kappaB kinase/NF-kappaB signaling | 2.83E-06 | 5.924981151 | 2.25413403 | 12 | 185 |
| GO:0043393 | regulation of protein binding | 3.35E-06 | 5.822875494 | 2.290687554 | 12 | 188 |
| GO:0070509 | calcium ion import | 5.77E-06 | 9.348699764 | 0.974760661 | 8 | 80 |
| GO:0046718 | viral entry into host cell | 8.88E-06 | 6.467618813 | 1.718015666 | 10 | 141 |
| GO:0030308 | negative regulation of cell growth | 1.18E-05 | 5.598090306 | 2.16884247 | 11 | 178 |
| GO:0002720 | positive regulation of cytokine production involved in immune response | 1.68E-05 | 9.771604938 | 0.816362054 | 7 | 67 |
| GO:0043491 | protein kinase B signaling | 1.78E-05 | 5.339459459 | 2.266318538 | 11 | 186 |

## 3 Discussion

In conclusion, G3DC utilizes both gene expression levels and gene-gene interaction graph, resulting in both clustering results that were faithful to cell types, as well as pointing to genes and pathways that differentiate the clusters. The regularization term of local structure reconstruction prevents the encoder model from overfitting and ensures the final clustering performance. Compared with several state-of-the-art methods, G3DC has proven its high accuracy and outstanding performance in data embedding. In particular, G3DC does not require extensive data preprocessing, other than scaling the gene expression vectors to ensure fair gene selection, since the feature importance can be learned through optimization.

As scRNA-seq technology gradually becomes a revolutionary tool in biomedical applications, not only clustering performance but also biological interpretation is required for further functional understanding. G3DC is able to provide meaningful interpretations based on gene importance estimated by the model. In our tested data, we selected the top significant genes whose first layer weights in the network were the highest. With the trace regularization optimization term, G3DC encourages adjacent genes to share similar weights. The method tended to select genes that are on sub-networks. Given the network encapsules the functional relations between the genes, such selection results greatly facilitates interpretation, and supports the discovery of biological functions based on single-cell level analysis.

## 4 Methods

G3DC aims to select informative and discriminative features from the high-dimensional zero-inflation scRNA-seq data for clustering cells while exploiting the intrinsic functional relations between genes. G3DC is based on a deep autoencoder, with four loss components involved. The first one is the reconstruction loss which helps discover the latent structure of the data and reduce redundancy. The second component is the KL divergence clustering loss, which measures the similarity between the proposed distribution and the pseudo-target distribution. The third component is the *ℓ*_2,1_-norm regularization that is able to identify the discriminative genes for clustering. The last component is graph regularization term Tr(*W*_1_*LW*_1_^⊤^), which lets adjacent genes share similar weights. This section details the architecture and optimization of G3DC.

### 4.1 Deep embedded clustering with gene graph information and feature selection

#### 4.1.1 Model initialization and reconstruction loss

The gene expression matrix *X* is a *p* ×*n* matrix with *p* representing the number of genes and *n* representing the number of cells. To cluster the *n* cells into *k* clusters, we use an encoder as a nonlinear mapping *f*_*θ*_ : ℝ^*p*^ →ℝ^*d*^ (*d*≪ *p*) to encode the columns of *X* to low-dimensional representations, which are then decoded by a decoder *g*_*θ*_*′* : ℝ^*d*^ →ℝ^*p*^ to reconstruct the columns of *X*, where *θ* and *θ*^′^ denote the parameters of the encoder network and decoder network respectively. More formally,

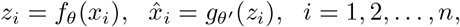

where *x*_*i*_ denotes the *i*-th column of *X*. The following reconstruction loss is considered in our G3DC

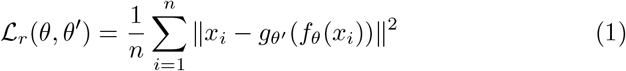

where 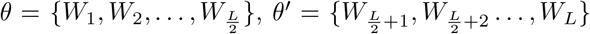, and *W*_*l*_ denotes the weight matrix of layer *l* of the deep autoencoder, *l* = 1, 2, …, *L*. By mapping the gene expression data to a low-dimensional space, the important information in the expression matrix is retained while the trivial part is discarded. Also, it lowers the computation complexity and facilitates the later clustering task.

Once the autoencoder is learned, K-means clustering is performed on the latent representations, i.e., *z*_1_, *z*_2_, …, *z*_*n*_, to obtain *k* initial cluster centers *μ*_1_, *μ*_2_, …, *μ*_*k*_. The *k* centers will be used and refined in a joint optimization stage detailed in Section 4.2.

#### 4.1.2 Soft label refinement and clustering module

Given the *k* cluster centers, a set of soft labels, denoted by *q*_*ij*_ can be obtained via using the Student’s t-distribution kernel to measure the similarity between embedded point *z*_*i*_ for cell *i* and *μ*_*j*_:

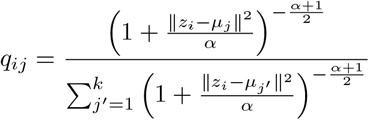

where *α* is the degree of freedom and we set it to be 1. *q*_*ij*_ can be regarded as the probability that data point *i* is assigned to cluster *j*. Since *q*_*ij*_ are not necessarily accurate, following [41], it could be refined as

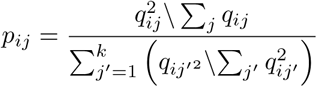

*P* = {*p*_*ij*_} is a target distribution and *Q* = {*q*_*ij*_} is a distribution learned by the neural network. It is expected that *Q* is as close as or even equal to *P*. Therefore, following [41], we propose to minimize the Kullback-Leibler (KL) divergence between *P* and *Q*, which yields the following loss

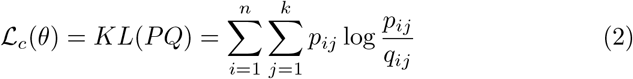

The KL divergence measures the difference between two distributions and *KL*(*PQ*) = 0 when *P* = *Q*. The encoder *f*_*θ*_ is fine-tuned by minimizing *L*_*c*_ and this process is a kind of self-training.

#### 4.1.3 *ℓ*_21_-norm, feature selection, and weight decay

The gene expression matrix is not only high-dimensional but also zero-inflated. Most genes do not provide useful information for clustering. We propose to select important and discriminative genes for clustering using the *ℓ*_2,1_-norm regularization. Specifically, we minimize

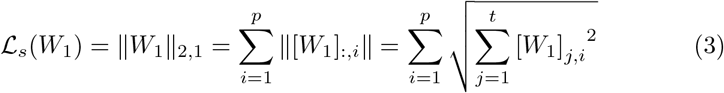

which makes some columns of *W*_1_ be zero. The genes corresponding to the nonzero columns of *W*_1_ are selected as significant genes.

#### 4.1.4 Gene-gene interaction and Laplacian regularization

The graph between genes contains information about the functional relationship between genes, which may be valuable in clustering. For the clustering results to be functionally meaningful, we expect adjacent genes in the graph to share similar weights. We extract the gene-gene relations from the HINT database [33] and construct the adjacency matrix of genes and denote it as *A*, where *A*_*ij*_ = 1 when gene *i* and gene *j* are functionally associated, otherwise *A*_*ij*_ = 0. The degree matrix is defined as *D* = diag(*d*_1_, *d*_2_, …, *d*_*p*_), where 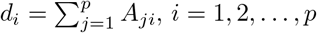. The symmetric normalized Laplacian matrix is defined as *L* = *I* −*D*^−1*/*2^*AD*^−1*/*2^, where *I* is an identity matrix of size *p*× *p*. To encourage those adjacent genes in the interaction graph to have similar weights, we use the following Laplacian regularization

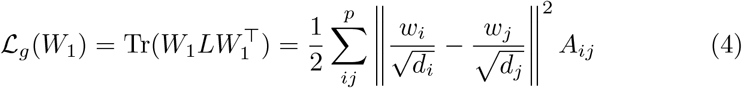

where *W*_1_ is the weight matrix of the first layer of the neural network, *w*_*i*_ denotes the *i*-th row of *W*_1_, and *tr* denotes the trace operator on a matrix. Minimizing *L*_*g*_(*W*_1_) will let 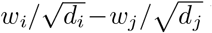 be small if *A*_*ij*_ = 1, which means the interacting genes share similar weights in the network. On the contrary, if *A*_*ij*_ = 0, we don’t impose any constraints.

### 4.2 Optimization

With the reconstruction loss (1), KL divergence (2), *ℓ*_21_-norm regularization (3), and graph Laplacian regularization (4), our final objective function is

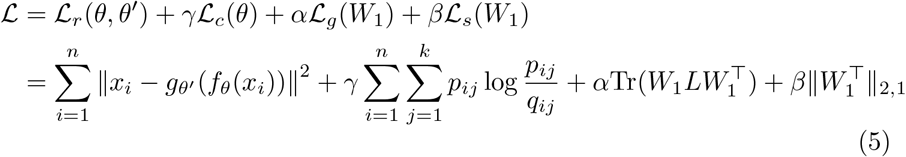

where *α, β* and *γ* are adjustable hyperparameters. Minimizing ℒ exploits the gene-gene interaction information, selects informative and discriminative genes, and clusters cells simultaneously.

We can use any gradient-based optimizer such as stochastic gradient descent or Adam [42] to minimize ℒ. The gradients of ℒ_*c*_ with respect to *z*_*i*_ and *μ*_*j*_ are as follows:

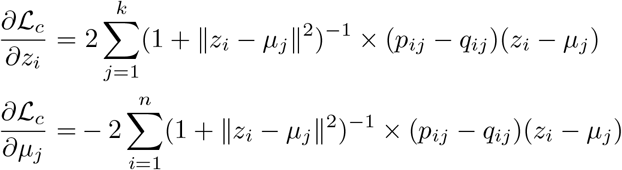

Now using the chain rule, we obtain the gradient of ℒ_*c*_ with respect to the parameters of the encoder, i.e.,

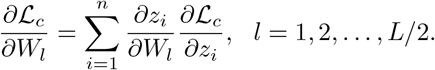

In addition, we have

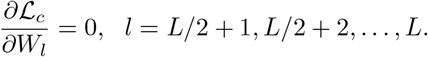

For convenience, let ℒ_*gs*_ = *αℒ*_*g*_ + *βℒ*_*s*_, which is only related to *W*_1_. The gradient of ℒ_*gs*_ with respect to *W*_1_ is

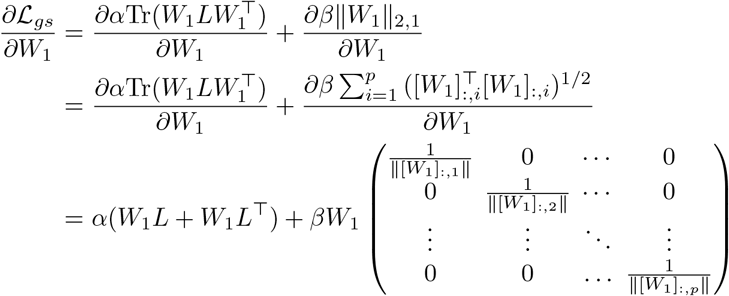

where [*W*_1_]_:*i*_ denotes the *i*-th column of *W*_1_. We have

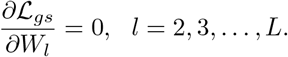

Now the gradients of ℒ with respect to the weight matrices are

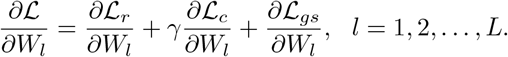

The gradient of ℒ with respect to the cluster centers are

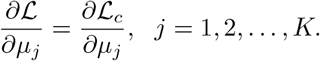

Then 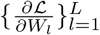 and 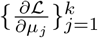 are used for the gradient-based optimizers. We have omitted the gradients of ℒ with respect to the bias terms for simplicity, though they are included in the actual computation.

The encoder part and decoder part are first pre-trained by the optimizer Adam [42], setting the initial learning rate *lr*=1e-3, weight decay = 1e-4. The optimizer for the clustering stage of Usoskin and Xin pancreas data is SGD with learning rate *lr*=1e-4, *momentum*=0.99 and weight decay = 1e-4. For Segerstolpe pancreas, Baron pancreas, and GSE96583 data, Adam optimizer is carried out in the clustering stage and the model is trained with learning rate *lr*=1e-4 and weight decay = 1e-5.

### 4.3 Evaluation metrics

To evaluate the clustering performance from different perspectives, three widely-used evaluation metrics, normalized mutual information (NMI), unsupervised clustering accuracy (CA), and adjusted random index (ARI), were considered. Larger values of those measurements indicate higher consistency between the predicted labels and the true labels.

#### 4.3.1 Normalized Mutual Information (NMI)

NMI is defined as

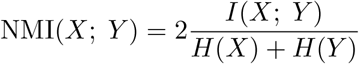

where *I*(*X*; *Y*) indicates the amount of mutual information between *X* and *Y*

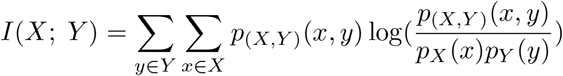

*H*(*X*) and *H*(*Y*) are the entropies that quantify uncertainties of *X* and *Y*

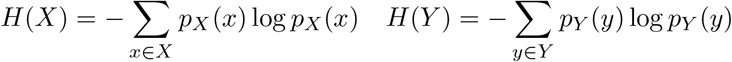

#### 4.3.2 Clustering Accuracy (CA)

The CA metric is commonly used to measure the difference between the predicted labels and the real labels in the clustering tasks. It tries to find the best matching between a cluster assignment of unsupervised algorithm and the ground truth assignment.

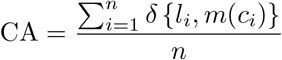

where *l*_*i*_ is the ground-truth and *c*_*j*_ is the cluster assignment using an unsupervised clustering algorithm. *δ*(*x, x*) is an indicator function that if *x* = *y, δ*(*x, y*) = 1, otherwise, *δ*(*x, y*) = 0. Here we choose the Hungarian algorithm to optimize the assignment between the predicted labels and the true labels.

#### 4.3.3 Adjusted Rank Index (ARI)

ARI is an improved version of Rand Index (RI). It is computed based on the contingency table between cluster memberships and true class labels as follows,

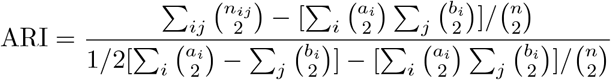

where *n*_*ij*_ is the value obtained from contingency table, *a*_*i*_ is the sum of the *i*_*th*_ row of contingency table, and *b*_*j*_ is the sum of the *j*_*th*_ column of contingency table.

### 4.4 Selection of influential genes

In order to find which genes and biological processes contribute most to the differentiation of cell groups, we define the importance of genes based on their first-layer weights. For each of the *p* genes, the importance score (IS) is summarized from the first-layer weight *W*_1_ on the *t* hidden units:

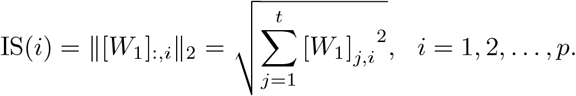

Genes receiving the highest importance scores are selected, and their functions further analyzed by standard functional analysis approaches.

### 4.5 Real datasets

Our method needs two inputs: 1) a gene expression matrix (*n*× *p*) with rows representing the cells and columns representing the genes. 2) a gene adjacent matrix (*p*× *p*) with each entry taking values 0 or 1, where 1 represents interaction. Five single-cell RNA-seq datasets (detailed below) were used to benchmark the performance of our method. Among them, one is a mouse dataset and the other four are human datasets. Table 1 shows the summary of the datasets.

#### 4.5.1 Segerstolpe pancreas dataset

The Segerstolpe pancreas dataset is an scRNA-seq dataset of human pancreas islets [43]. We used a sub-dataset from wuhaolab.org that consists 1000 highly variable genes and 702 cells. There are 6 cell types, including alpha, beta, gamma, ductal, delta, and endothelial cells. A total of 1391 interactions are included in the HINT database between the 1000 genes.

#### 4.5.2 Baron pancreas data

The Baron pancreas dataset is a single-cell transcriptome of the human pancreas [44]. We downloaded the sub-dataset from wuhaolab.org that contains the measurement of 2000 highly variable genes in 2812 cells. There are 6 cell types in this dataset, including alpha, beta, gamma, ductal, delta, and endothelial cells. A total of 5441 interactions are included in the HINT database between the 2000 genes.

#### 4.5.3 Xin pancreas data

The Xin pancreas data is an RNA sequencing of single human islet cells with the accession GSE81608 [45]. This dataset contains a hundred thousand of genes and 2000 highly variable genes were selected. There are 1600 cells in this dataset. The 4 cell types include alpha, beta, gamma, and delta cells. A total of 2201 interactions are included in the HINT database between the 2000 genes.

#### 4.5.4 GSE96583 data

The GSE96583 data is a droplet-based single-cell RNA-sequencing data on peripheral blood mononuclear cells (PBMC) [46].The original dataset is obtained from GEO with accession number GSE96583. There are 2180 cells belonging to 8 cell types including CD14+ monocytes, CD4 T cells, CD8 T cells, NK cells, FCGR3A+ monocytes, B cells, dendritic cells, and megakaryocytes. After deleting genes with 0 expression levels across all cells, 14141 genes were left, and we use them to cluster the 2180 cells. A total of 62782 interactions are included in the HINT database between the 14141 genes.

#### 4.5.5 Usoskin data

The Usoskin single-cell RNAseq data was used to analyze sensory neuron cells. This data was generated from the mouse lumbar dorsal root ganglion [47]. The original dataset can be retrieved from GEO database with accession number GSE59739. There are in total 25334 genes and 610 cells with 4 cell types, including non-peptidergic nociceptor cells (NP), peptidergic nociceptor cells (PEP), neurofilament containing cells (NF), and tyrosine hydroxylase containing cells (TH). A total of 83175 interactions are included in the HINT database between the genes.

## 5 Data and code availability

The source code, trained model weights and the real scRNA-seq dataset used for experiments are available in Github: https://github.com/Ketherine0/G3DC.

## Acknowledgements

This work was partially supported by Shenzhen Research Institute of Big Data, and the University Development Fund of CUHK-Shenzhen.

## References

[1] Hartigan, J.A., Wong, M.A.: Algorithm as 136: A k-means clustering algorithm. Journal of the royal statistical society. series c (applied statistics) 2

[2] Yau, C., et al.: pcareduce: hierarchical clustering of single cell transcriptional profiles. BMC bioinformatics 17(1), 1–11 (2016)

[3] Jolliffe, I.T.: Principal Component Analysis for Special Types of Data. Springer, ??? (2002)

[4] Herman, J.S., Grün, D., et al.: Fateid infers cell fate bias in multipotent progenitors from single-cell rna-seq data. Nature methods 15(5), 379–386 (2018)

[5] Hicks, S.C., Liu, R., Ni, Y., Purdom, E., Risso, D.: mbkmeans: Fast clustering for single cell data using mini-batch k-means. PLoS computational biology 17(1), 1008625 (2021)

[6] Lin, P., Troup, M., Ho, J.W.: Cidr: Ultrafast and accurate clustering through imputation for single-cell rna-seq data. Genome biology 18(1), 1–11 (2017)

[7] Zeisel, A., Muñoz-Manchado, A.B., Codeluppi, S., Lönnerberg, P., La Manno, G., Juréus, A., Marques, S., Munguba, H., He, L., Betsholtz, C., et al.: Cell types in the mouse cortex and hippocampus revealed by single-cell rna-seq. Science 347(6226), 1138–1142 (2015)

[8] Guo, M., Wang, H., Potter, S.S., Whitsett, J.A., Xu, Y.: Sincera: a pipeline for single-cell rna-seq profiling analysis. PLoS computational biology 11(11), 1004575 (2015)

[9] Ng, A., Jordan, M., Weiss, Y.: On spectral clustering: Analysis and an algorithm. Advances in neural information processing systems 14 (2001)

[10] Shi, J., Malik, J.: Normalized cuts and image segmentation. IEEE Transactions on pattern analysis and machine intelligence 22(8), 888–905 (2000)

[11] Blondel, V.D., Guillaume, J.-L., Lambiotte, R., Lefebvre, E.: Fast unfolding of communities in large networks. Journal of statistical mechanics: theory and experiment 2008(10), 10008 (2008)

[12] Traag, V.A., Waltman, L., Van Eck, N.J.: From louvain to leiden: guaranteeing well-connected communities. Scientific reports 9(1), 1–12 (2019)

[13] Satija, R., Farrell, J.A., Gennert, D., Schier, A.F., Regev, A.: Spatial reconstruction of single-cell gene expression data. Nature biotechnology 33(5), 495–502 (2015)

[14] Wolf, F.A., Angerer, P., Theis, F.J.: Scanpy: large-scale single-cell gene expression data analysis. Genome biology 19(1), 1–5 (2018)

[15] Anuar, S.H.H., Abas, Z.A., Yunos, N.M., Zaki, N.H.M., Hashim, N.A., Mokhtar, M.F., Asmai, S.A., Abidin, Z.Z., Nizam, A.F.: Comparison between louvain and leiden algorithm for network structure: A review. In: Journal of Physics: Conference Series, vol. 2129, p. 012028 (2021). IOP Publishing

[16] Xu, C., Su, Z.: Identification of cell types from single-cell transcriptomes using a novel clustering method. Bioinformatics 31(12), 1974–1980 (2015)

[17] Wang, B., Ramazzotti, D., De Sano, L., Zhu, J., Pierson, E., Batzoglou, S.: Simlr: A tool for large-scale genomic analyses by multi-kernel learning. Proteomics 18(2), 1700232 (2018)

[18] Qiu, X., Mao, Q., Tang, Y., Wang, L., Chawla, R., Pliner, H.A., Trapnell, C.: Reversed graph embedding resolves complex single-cell trajectories. Nature methods 14(10), 979–982 (2017)

[19] Yang, Y., Huh, R., Culpepper, H.W., Lin, Y., Love, M.I., Li, Y.: Safe-clustering: single-cell aggregated (from ensemble) clustering for single-cell rna-seq data. Bioinformatics 35(8), 1269–1277 (2019)

[20] Huh, R., Yang, Y., Jiang, Y., Shen, Y., Li, Y.: Same-clustering: Singlecell a ggregated clustering via m ixture model e nsemble. Nucleic acids research 48(1), 86–95 (2020)

[21] Zhu, X., Li, J., Li, H.-D., Xie, M., Wang, J.: Sc-gpe: A graph partitioning-based cluster ensemble method for single-cell. Frontiers in Genetics 11, 604790 (2020)

[22] Ranjan, B., Schmidt, F., Sun, W., Park, J., Honardoost, M.A., Tan, J., Arul Rayan, N., Prabhakar, S.: scconsensus: combining supervised and unsupervised clustering for cell type identification in single-cell rna sequencing data. BMC bioinformatics 22(1), 1–15 (2021)

[23] Zhu, X., Zhang, J., Xu, Y., Wang, J., Peng, X., Li, H.-D.: Singlecell clustering based on shared nearest neighbor and graph partitioning. Interdisciplinary Sciences: Computational Life Sciences 12(2), 117–130 (2020)

[24] Zhu, X., Li, H.-D., Guo, L., Wu, F.-X., Wang, J.: Analysis of singlecell rna-seq data by clustering approaches. Current Bioinformatics 14(4), 314–322 (2019)

[25] Kiselev, V.Y., Kirschner, K., Schaub, M.T., Andrews, T., Yiu, A., Chandra, T., Natarajan, K.N., Reik, W., Barahona, M., Green, A.R., et al.: Sc3: consensus clustering of single-cell rna-seq data. Nature methods 14(5), 483–486 (2017)

[26] Eraslan, G., Simon, L.M., Mircea, M., Mueller, N.S., Theis, F.J.: Single-cell rna-seq denoising using a deep count autoencoder. Nature communications 10(1), 1–14 (2019)

[27] Tian, T., Wan, J., Song, Q., Wei, Z.: Clustering single-cell rna-seq data with a model-based deep learning approach. Nature Machine Intelligence 1(4), 191–198 (2019)

[28] Li, X., Wang, K., Lyu, Y., Pan, H., Zhang, J., Stambolian, D., Susztak, K., Reilly, M.P., Hu, G., Li, M.: Deep learning enables accurate clustering with batch effect removal in single-cell rna-seq analysis. Nature communications 11(1), 1–14 (2020)

[29] Guo, X., Gao, L., Liu, X., Yin, J.: Improved deep embedded clustering with local structure preservation. In: Ijcai, pp. 1753–1759 (2017)

[30] Tran, B., Tran, D., Nguyen, H., Ro, S., Nguyen, T.: sccan: single-cell clustering using autoencoder and network fusion. Scientific reports 12(1), 1–10 (2022)

[31] Wang, T., Li, B., Nabavi, S.: Single-cell rna sequencing data clustering using graph convolutional networks. In: 2021 IEEE International Conference on Bioinformatics and Biomedicine (BIBM), pp. 2163–2170 (2021). IEEE

[32] Kipf, T.N., Welling, M.: Semi-supervised classification with graph convolutional networks. arXiv preprint arXiv:1609.02907 (2016)

[33] Patil, A., Nakamura, H.: Hint: a database of annotated protein-protein interactions and their homologs. Biophysics 1, 21–24 (2005)

[34] Van der Maaten, L., Hinton, G.: Visualizing data using t-sne. Journal of machine learning research 9(11) (2008)

[35] Beissbarth, T., Speed, T.P.: Gostat: find statistically overrepresented gene ontologies within a group of genes. Bioinformatics 20(9), 1464–1465 (2004)

[36] Lin, R., Bao, X., Wang, H., Zhu, S., Liu, Z., Chen, Q., Ai, K., Shi, B.: Trpm2 promotes pancreatic cancer by pkc/mapk pathway. Cell Death Dis 12(6), 585 (2021). https://doi.org/10.1038/s41419-021-03856-9

[37] Khoo, S., Gibson, T.B., Arnette, D., Lawrence, M., January, B., McGlynn, K., Vanderbilt, C.A., Griffen, S.C., German, M.S., Cobb, M.H.: Map kinases and their roles in pancreatic beta-cells. Cell Biochem Biophys 40(3 Suppl), 191–200 (2004). https://doi.org/10.1385/cbb:40:3:191

[38] Fasolino, M., Schwartz, G.W., Patil, A.R., Mongia, A., Golson, M.L., Wang, Y.J., Morgan, A., Liu, C., Schug, J., Liu, J., Wu, M., Traum, D., Kondo, A., May, C.L., Goldman, N., Wang, W., Feldman, M., Moore, J.H., Japp, A.S., Betts, M.R., Consortium, H., Faryabi, R.B., Naji, A., Kaestner, K.H., Vahedi, G.: Single-cell multi-omics analysis of human pancreatic islets reveals novel cellular states in type 1 diabetes. Nat Metab 4(2), 284–299 (2022). https://doi.org/10.1038/s42255-022-00531-x

[39] He, X., Gao, F., Hou, J., Li, T., Tan, J., Wang, C., Liu, X., Wang, M., Liu, H., Chen, Y., Yu, Z., Yang, M.: Metformin inhibits mapk signaling and rescues pancreatic aquaporin 7 expression to induce insulin secretion in type 2 diabetes mellitus. J Biol Chem 297(2), 101002 (2021). https://doi.org/10.1016/j.jbc.2021.101002

[40] Bogdan, C.: Nitric oxide and the immune response. Nat Immunol 2(10), 907–16 (2001). https://doi.org/10.1038/ni1001-907

[41] Xie, J., Girshick, R., Farhadi, A.: Unsupervised deep embedding for clustering analysis. In: International Conference on Machine Learning, pp. 478–487 (2016). PMLR

[42] Kingma, D.P., Ba, J.: Adam: A method for stochastic optimization. arXiv preprint arXiv:1412.6980 (2014)

[43] Segerstolpe, Å., Palasantza, A., Eliasson, P., Andersson, E.-M., Andréasson, A.-C., Sun, X., Picelli, S., Sabirsh, A., Clausen, M., Bjursell, M.K., et al.: Single-cell transcriptome profiling of human pancreatic islets in health and type 2 diabetes. Cell metabolism 24(4), 593–607 (2016)

[44] Baron, M., Veres, A., Wolock, S.L., Faust, A.L., Gaujoux, R., Vetere, A., Ryu, J.H., Wagner, B.K., Shen-Orr, S.S., Klein, A.M., et al.: A single-cell transcriptomic map of the human and mouse pancreas reveals inter-and intra-cell population structure. Cell systems 3(4), 346–360 (2016)

[45] Xin, Y., Kim, J., Okamoto, H., Ni, M., Wei, Y., Adler, C., Murphy, A.J., Yancopoulos, G.D., Lin, C., Gromada, J.: Rna sequencing of single human islet cells reveals type 2 diabetes genes. Cell metabolism 24(4), 608–615 (2016)

[46] Kang, H.M., Subramaniam, M., Targ, S., Nguyen, M., Maliskova, L., McCarthy, E., Wan, E., Wong, S., Byrnes, L., Lanata, C.M., et al.: Multiplexed droplet single-cell rna-sequencing using natural genetic variation. Nature biotechnology 36(1), 89–94 (2018)

[47] Usoskin, D., Furlan, A., Islam, S., Abdo, H., Lönnerberg, P., Lou, D., Hjerling-Leffler, J., Haeggström, J., Kharchenko, O., Kharchenko, P.V., et al.: Unbiased classification of sensory neuron types by large-scale singlecell rna sequencing. Nature neuroscience 18(1), 145–153 (2015)

